# Candidate Genes Underlying *Wolbachia*-Associated Plastic Recombination Revealed by Ovarian Transcriptomics of *D. melanogaster*

**DOI:** 10.1101/2022.04.28.489783

**Authors:** Sophia I. Frantz, Clayton M. Small, William A. Cresko, Nadia D. Singh

**Affiliations:** Institute of Ecology and Evolution, University of Oregon, Eugene, OR

**Keywords:** *Wolbachia*, transcriptomics, host-microbe interactions, ovary, plasticity, *Drosophila melanogaster*, *Wolbachia pipientis*, recombination

## Abstract

Phenotypic plasticity is prevalent in nature and can facilitate the acclimation of organisms to changing environments. Recombination rate is plastic in a diversity of organisms and under a variety of stressful conditions. However, the recent finding that *Wolbachia pipientis* induces plastic recombination in *Drosophila melanogaster* was surprising because *Wolbachia* is not strictly considered a stressor to this host. We investigate the molecular mechanisms of *Wolbachia*-associated plastic recombination by comparing the ovarian transcriptomes of *D. melanogaster* infected and uninfected with *Wolbachia*. Our data suggest that infection explains a small amount of transcriptional variation but specifically affects genes related to cell cycle, translation, and metabolism. We also find enrichment of cell division and recombination processes among genes with infection-associated differential expression. Broadly, the transcriptomic changes identified in this study provide insight for the mechanisms of microbe-mediated plastic recombination, an important but poorly understood facet of host-microbe dynamics.

**Significance Statement:** Though it is documented that *Wolbachia* is associated with increased recombination *in D. melanogaster*, the underlying mechanisms remain unknown. Here we use ovarian transcriptomics in *Wolbachia*-infected and uninfected flies to identify candidate genes underlying *Wolbachia*-associated plastic recombination. We find that infection alters ovarian gene expression in subtle ways, and also identify exciting candidate genes for functional analysis in subsequent work. These candidate genes may be the first step in determining the molecular mechanisms underlying *Wolbachia*-associated plastic recombination. Moreover, our data contribute to the growing body of knowledge surrounding how *Wolbachia* affects host gene expression, and highlights how context-dependent these effects are.

## Introduction

Under a changing or stressful environment, populations may adapt over generations, and, more immediately, individuals in these populations may acclimate within a lifetime via phenotypic plasticity. Phenotypic plasticity is the phenomenon by which one genotype produces multiple phenotypes in response to different environmental conditions (Bradshaw 1965). Phenotypic plasticity is pervasive in nature and includes well-studied examples such as seasonal color changes of *Precis octavia* butterflies (Brakefield 1987) and predator-induced morphological changes in *Daphnia* (Tollrian 1995). Triggering conditions may be exogenous— such as temperature (Scheepens, et al. 2018), or endogenous— such as microbial presence (Verbon and Liberman 2016).

Although many previously studied cases of phenotypic plasticity have involved morphological, behavioral or life history traits, genetic recombination itself can also be plastic (for review see Modliszewski and Copenhaver 2017; Stapley, et al. 2017). Plastic recombination has been documented in diverse organisms and under a variety of environmental conditions, such as in yeast under osmotic stress (Gao, et al. 2008), in mice under social stress (Belyaev and Borodin 1982), and in humans with increasing maternal age (Hussin, et al. 2011). The mechanistic link between environmental stress and plastic recombination is largely unknown, though studies in yeast have identified known stress response genes that also regulate recombination rate at specific genomic locations (Gao, et al. 2008; Kon, et al. 1998). Plastic recombination is a major contributor to variation in recombination rate (Agrawal, et al. 2005), and its contribution to genetic diversity overall requires understanding of the conditions in which it manifests.

The study of recombination in the genetic model organism *Drosophila melanogaster* led to key insights about conditions that trigger plastic recombination, which occurred in response to extreme temperature (Grell 1978; Plough 1917, 1921), starvation (Neel 1941), and age (Hayman and Parsons 1961; Stern 1926). Recent work also supports the effect of age (Hunter, et al. 2016b) and temperature on plastic recombination (Jackson, et al. 2015; Kohl and Singh 2018), in addition to other environmental cues such as desiccation (Aggarwal, et al. 2019) and parasitic infection (Andronic 2012; Singh, et al. 2015; Zilio, et al. 2018).

The recent finding that the *w*Mel strain of the endosymbiont *Wolbachia pipientis* also induces plastic recombination in *Drosophila melanogaster* (Bryant and Newton 2020; Singh 2019) was surprising in the context of these previous studies of plastic recombination, since *w*Mel is not generally considered a stressor. Strains of *D. melanogaster* naturally infected with the *w*Mel strain of *Wolbachia* produced a higher proportion of recombinant offspring than individuals of the same strains cleared of *w*Mel. Recombination was also shown to increase at an X-chromosome locus but not at a third chromosome locus, which suggests that *Wolbachia* affects recombination at specific regions but not globally across the genome (Singh 2019).

Although never previously associated with plastic recombination, *Wolbachia* is a well-studied endosymbiont that resides in the ovaries of its host and is maternally transmitted. It is estimated that up to 40-60% of arthropod species are infected with a species of the *Wolbachia* genus (Hilgenboecker, et al. 2008; Zug and Hammerstein 2012). The proportion of infected individuals varies by population, with some populations exhibiting very high frequencies of infection (Kriesner, et al. 2013; Weeks, et al. 2007){Kaur, 2021 #171}.

*Wolbachia* exerts a wide range of effects on their various hosts. That is, Wolbachia is a bacterium that employs a range of parasitic and mutualistic traits, sometimes even from the same strain. In some host species, *Wolbachia* infection is classified as harmful, causing reduced reproductive output through four main mechanisms: cytoplasmic incompatibility (CI), male-killing, feminization, and parthenogenesis (for review see Werren, et al. 2008). These mechanisms often ensure *Wolbachia*’s propagation to the next generation. Theoretical and empirical evidence also suggests these parasitic interactions can evolve to become more mutualistic over time (Ballard 2004; Weeks, et al. 2007). In other cases, *Wolbachia* can confer advantages such as increased reproductive output and protection from viral infection (Vavre, et al. 1999). Some hosts receive essential nutrients from *Wolbachia* (Brownlie, et al. 2009; Hosokawa, et al. 2010), and *Drosophila paulistorum* even presents an extreme example of obligate mutualism in which *Wolbachia* is needed for gonad development (Miller, et al. 2010).

The effect of *Wolbachia* therefore strongly depends on the particular host species and genotype as well as the strain of *Wolbachia*. In its *D. melanogaster* host, *w*Mel induces only weak CI in laboratory strains and even weaker CI in the wild (Hoffmann, et al. 1998). Therefore, the high global infection rate of *w*Mel (estimated 34% in wild populations of *D. melanogaster*) cannot be explained by reproductive manipulation through CI alone and may instead be explained by benefits conferred to the host (Serga, et al. 2014; Solignac, et al. 1994). Benefits such as increased survival and fecundity have been documented in *w*Mel -infected strains of *D. melanogaster* (Fry, et al. 2004; Fry and Rand 2002). In fact, the same strains that exhibit increased recombination with *Wolbachia* infection also exhibited increased reproductive output (Singh 2019).

Despite this evidence that *w*Mel is beneficial to its *D. melanogaster* host, *w*Mel also induces increased recombination (Bryant and Newton 2020; Singh 2019), a trait sometimes associated with stressful conditions. Interestingly, flies infected with the virulent *Wolbachia* strain *w*MelPop show an exaggerated response (Bryant and Newton 2020). This novel mediator of plastic recombination and its ambiguous relationship to stress prompted us to investigate the molecular mechanisms that mediate *Wolbachia*-associated plastic recombination. We hypothesized that the molecular genetic mechanism underlying increased recombination in *Wolbachia*-infected flies would be reflected in host transcriptional changes. To test this hypothesis we performed RNA-sequencing and differential expression analysis to compare the transcriptomes of *Wolbachia*-uninfected and infected flies. Since the ovaries are the site of both *Wolbachia* colonization and meiotic recombination, we sequenced the ovarian transcriptome.

We focused primarily on characterizing global ovarian transcriptome changes and identifying candidate genes that mediate *Wolbachia*-associated plastic recombination. The use of four different strains of *D. melanogaster* allowed us to disentangle transcriptional variation due to infection, genotype, and the interaction of infection and genotype. We note that all four strains of *D. melanogaster* are infected with *w*Mel haplotype I. Though there are certainly genetic differences among the *w*Mel halplotype I strains, these differences are limited and levels of genetic variation within these *Wolbachia* strains is far lower than the genetic variation between fly strains (Richardson, et al. 2012). Moreover, all four strains of *D. melanogaster* have the type I mitochondrial haplotype {Richardson, 2012 #74}. Polymorphism within the Drosophila Genetic Reference Panel {Mackay, 2012 #162} mitotypes (from which these four strains were chosen) is five times lower than polymorphism within the nuclear genome {Richardson, 2012 #74}. Collectively, these data indicate that the effects of variation within Wolbachia and within the mitochondrial genome are likely to be swamped by the effects of genotype variation between D. melanogaster strains. Thus, our experimental design allows us to focus on host genotype, infection status, and interaction effects. Our results suggest the contribution of *Wolbachia* to ovarian transcriptional variation is limited, variable, and mediated by the host genotype. Finally, we identified a subset of genes with functions related to cell division, translation, and metabolism that are differentially regulated under *Wolbachia* infection.

## Results

### Wolbachia’s influence on ovary transcriptional differences is variable while the influence of genotype is consistent across samples

To understand the relative extent to which transcriptional variation depends on *Wolbachia* infection status and genotype, we visualized this variation across samples using non-metric multidimensional scaling (nMDS) (Figure 1a). Arguably the strongest signal in the nMDS space is a batch effect. In our experimental design, we conducted the experiment at two different times. In the first experiment, we collected and sequenced flies for strains RAL73 and RAL 783. In the second experiment, we did the same for RAL306 and RAL856. The batch effect in the analysis therefore reflects the time at which the experiment was conducted, and this is reflected in the nMDS space. Specifically, the cluster of samples prepared in the first batch (RAL73 and RAL783) is distinct from the cluster of samples prepared in the second batch (RAL306 and RAL856) and indeed show stronger evidence for separation in nMDS space than any other grouping. Isolating this batch effect allows us to analyze biological patterns within each batch, which can be treated as two separate but parallel experiments.

**Figure 1.**
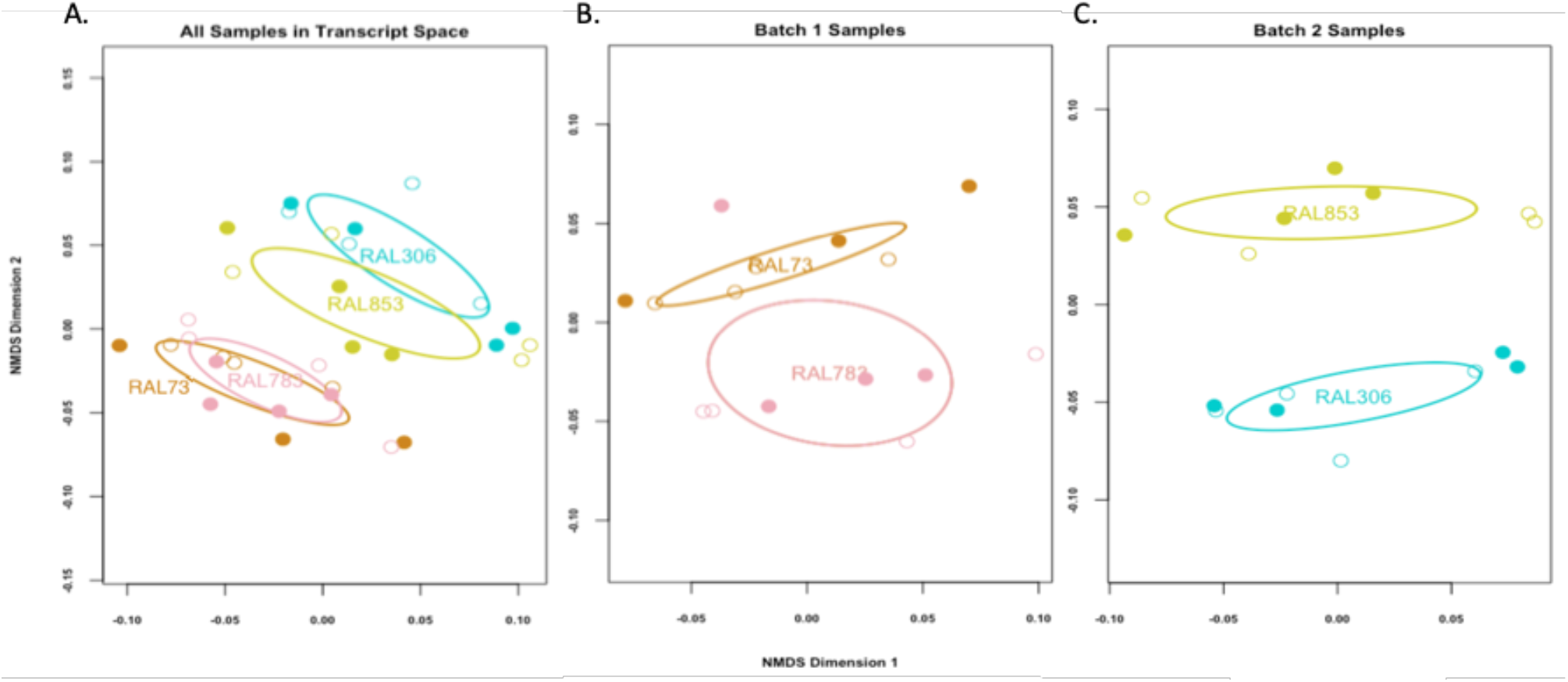
Global ovarian transcriptome nMDS plots. Each sample is represented in 2-dimensional transcript space with a circle, with open circles representing uninfected samples and filled circles representing infected samples. Each genotype is depicted with a different color. The ellipses mark the 95% confidence intervals of the centroid for the samples from each genotype. Note that nMDS dimensions do not quantify distances between points in the space, just their ranks. a) The samples cluster based on genotype and infection status, with genotype contributing to the most obvious clustering. The batch effect is also visible: the RAL73 and 783 samples form a cluster while the RAL306 and RAL853 samples form a separate cluster. b) Dissimilarity was calculated only between samples in batch 1. Samples cluster by genotype and subtly by infection status for RAL73. c) Dissimilarity was calculated only between samples in batch 2. The samples group by genotype and there is no obvious grouping by infection status.

The strongest biological pattern in this analysis is the grouping of samples based on genotype. Samples of the same genotype are often associated with each other regardless of infection status. Both genotype and infection status contribute to transcriptional differences among samples in batch 1 when plotting samples from each batch on a separate nMDS plot (Figure 1b). In contrast, for the batch 2 samples alone (Figure 1c) suggests that genotype is a strong determinant of transcriptional differences while infection status is not. Overall, the contribution of *Wolbachia* to global ovarian transcriptional variation is subtle but depends on genotype, while genotype consistently accounts for transcriptional differences in these samples.

To statistically test and quantify the patterns observed in the nMDS plot, we used permutational multivariate analysis of variance (perMANOVA) to estimate the percent of variation in gene expression explained by infection status, genotype, and the interaction of these factors (Table 1). Consistent with the nMDS patterns, genotype accounts for a large and statistically significant proportion of variation in both batch 1 (R^2^ = 0.17, *P* = 0.006) and batch 2 samples (R^2^ = 0.26, *P* = 0.01). For batch 1, infection status explains a similarly large percent of variation as does genotype (R^2^ = 0.19, *P* = 0.003). In batch 1, the interaction of genotype and infection also explains a statistically significant proportion of transcriptional variation (R^2^ = 0.13, *P* = 0.02). This interaction means *Wolbachia*’s contribution to transcriptional differences in these samples depends on host genotype (for RAL73 or RAL783).

**Table 1.**
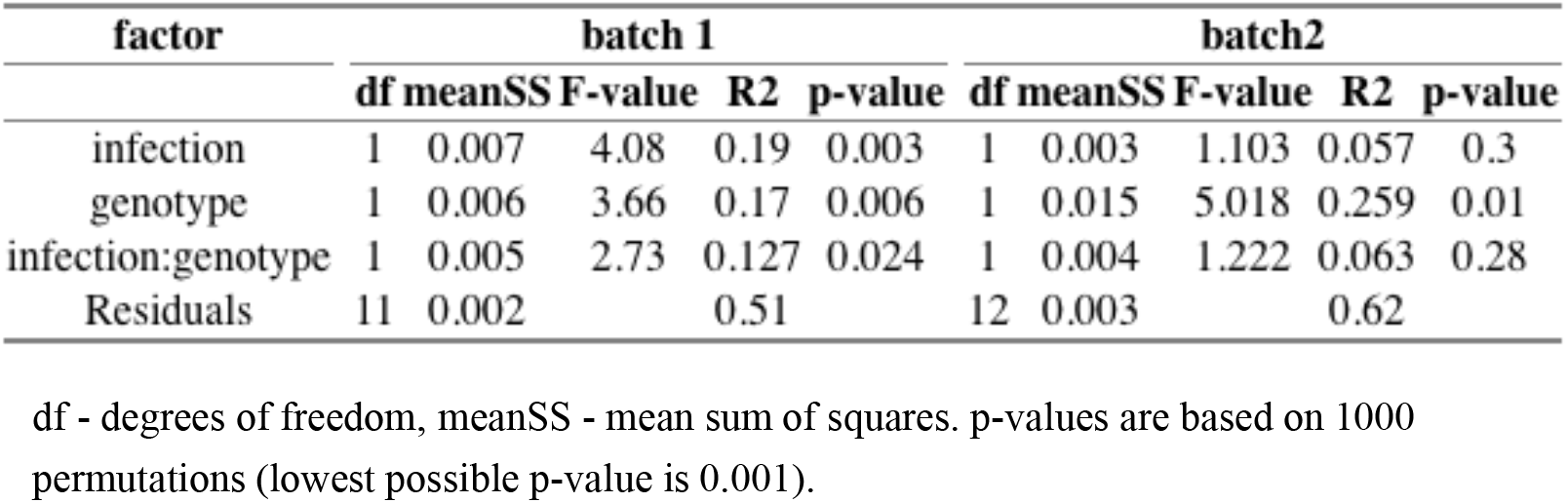
Summary of perMANOVA test results. Infection status and genotype explains a large and significant proportion of the dissimilarity among samples in batch 1 while genotype alone explains the largest proportion of accounted dissimilarity in batch 2. The perMANOVA is based on Bray-Curtis dissimilarities using gene expression differences across samples. The test was performed separately for each batch.

| factor | batch 1 |  |  |  |  | batch2 |  |  |  |  |
| --- | --- | --- | --- | --- | --- | --- | --- | --- | --- | --- |
|  | df | meanSS | F-value | R2 | p-value | df | meanSS | F-value | R2 | p-value |
| infection | 1 | 0.007 | 4.08 | 0.19 | 0.003 | 1 | 0.003 | 1.103 | 0.057 | 0.3 |
| genotype | 1 | 0.006 | 3.66 | 0.17 | 0.006 | 1 | 0.015 | 5.018 | 0.259 | 0.01 |
| infection:genotype | 1 | 0.005 | 2.73 | 0.127 | 0.024 | 1 | 0.004 | 1.222 | 0.063 | 0.28 |
| Residuals | 11 | 0.002 |  | 0.51 |  | 12 | 0.003 |  | 0.62 |  |

df - degrees of freedom, meanSS - mean sum of squares. p-values are based on 1000 permutations (lowest possible p-value is 0.001).

In contrast, infection status accounts for a smaller and non-significant proportion of transcriptional variation among samples in batch 2 (R^2^ = 0.06, *P* = 0.3). Consistent with the small effect of infection status, the interaction effect for the samples in batch 2 was also small and nonsignificant (R^2^ = 0.06, *P* = 0.28). In conclusion, genotype and infection status contribute about equally to the variation in transcription among the samples in batch 1, whereas genotype accounts for much more of the variation than does infection status in batch 2. The contribution of infection status to host gene expression varies, while the contribution of genotype is consistently large and significant in these samples.

### Wolbachia infection induces differential gene expression in D. melanogaster ovaries

The transcriptional profile of *Wolbachia* infection can also be described by the individual genes that are differentially regulated in *Wolbachia*-infected flies as compared to uninfected flies. In batch 1, 1104 genes (6% of all transcripts) are differentially expressed, with 618 upregulated in *Wolbachia*-infected flies and 486 downregulated. The differentially expressed genes, along with their fold change, direction of change, mean number of counts, test statistic, and adjusted p-values sorted in ascending order are shown in Supplementary Table 1. In batch 2, 96 genes (0.5% of all transcripts) are differentially expressed, with 36 upregulated and 60 downregulated genes (Supplementary Table 2). This difference between the number of genes identified in the two batches is consistent with the multivariate analysis, that infection status does not account for a large proportion of global transcriptional differences in batch 2. By comparing the sets of genes from the two batches we find only 26 genes that are consistently differentially expressed across these samples despite genotype and batch differences (Supplementary Table 3).

### Differentially expressed genes are randomly distributed across the genome

In the Drosophila early ovarian transcriptome, genomic regions with higher rates of transcription also have higher rates of recombination (Adrian and Comeron 2012). Since *Wolbachia* infection leads to increased recombination at certain genomic regions (Bryant and Newton 2020; Singh 2019), we tested whether infection may also be associated with altered levels of transcription at certain genomic regions. To do so, we performed runs test for randomness (Bradley 1968) to determine whether the binary sequence of genes that are categorized as differentially expressed or not is random across the genome. We performed the runs test for each chromosome arm and never rejected the null hypothesis of randomness, suggesting that the location of differentially expressed genes across the genome is random. This random placement of differentially expressed genes across each chromosome arm is illustrated in Figure 2. The visualization and runs test results provide no evidence that *Wolbachia* infection is associated with differential expression of genes at particular genomic locations.

**Figure 2.**
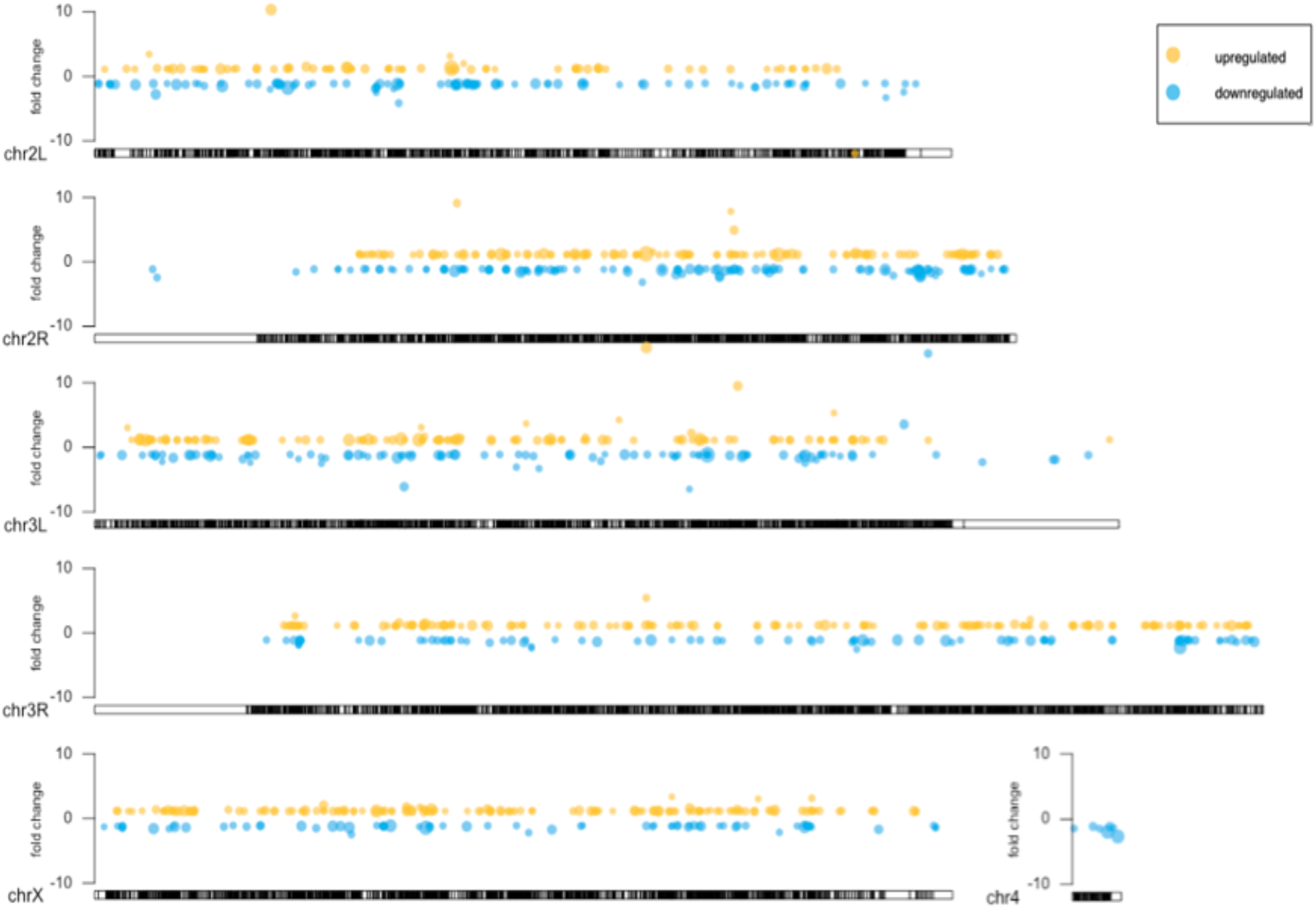
Location of differentially expressed genes from batch 1 across the genome for each chromosome. Differentially expressed genes are distributed randomly across the genome as determined by the runs test for randomness. Genes with an adjusted p-value < 0.05 are shown at their position on the chromosome, represented by the ideogram. Each circle represents a gene. The placement of the circle on the fold change axis shows the value of fold change difference when comparing uninfected and infected samples. The size of the circle is proportional to the range of p-values for all DE genes, with the biggest circles corresponding to the smallest p-values. Yellow represents upregulation and blue represents downregulation

### Differentially expressed genes are enriched in cell division processes

To better understand the function of these differentially expressed genes, we associated each gene with its most nested biological process GO terms (Mi, et al. 2010; Thomas, et al. 2003) and tested for overrepresentation of these terms. In batch 1, the 1104 differentially expressed genes contained significant overrepresentation of 99 GO terms. These GO terms are provided in Supplementary Table 5, sorted in order of highest to lowest enrichment score. The three most overrepresented terms are “spindle midzone assembly” (enrichment = 12.89, FDR = 0.008), “cytoplasmic translation” (enrichment = 8.56, FDR = 0.0001), and “mitotic spindle elongation (enrichment = 8.06, FDR = 0.027)”. Among the enriched terms were other cell cycle-related processes such as “centrosome separation” (enrichment = 6.78, FDR = 0.001) and “meiotic cytokinesis” (enrichment = 3.15, FDR = 0.033). Other common terms were those related to translation, such as “ribosomal small subunit assembly” (enrichment = 6.45, FDR = 0.001). Since batch 2 has only 96 differentially expressed genes, the statistical power to determine overrepresentation of GO terms was limited. Using a FDR threshold of 0.1, no GO terms were significantly enriched in batch 2. To get a sense of the genes processes we looked at the 55 GO terms with a raw *P*-value below 0.05 (Supplementary Table 6). The top terms in batch 2 are “N-acetylneuraminate catabolic process,”, “regulation of histone H4 acetylation involved in response to DNA damage stimulus,”, and “maintenance of rDNA” (enrichment > 100, *P* = 0.013 for all three terms). GO terms related to cell division processes are not among the most enriched in this small set of genes, but relevant terms include “synaptonemal complex assembly” (enrichment = 26.07, *P* = 0.04) and “female meiosis sister chromatid cohesion” (enrichment = 78.22, *P* = 0.019).

## Discussion

### Global transcription patterns suggest a consistent effect of genotype and variable influence of Wolbachia infection

Experiments from a wide range of model systems align with our finding that genotype significantly contributes to differences in global transcription (Hutter, et al. 2008; Poelstra, et al. 2014) and is consistent with evidence that variation in gene expression is heritable (Brem, et al. 2002; Dixon, et al. 2007). In fact, these same *D. melanogaster* strains (ours are a subset of the Drosophila Genetic Reference Panel) were used to quantify transcriptional differences due to genotype (Huang, et al. 2015). Of 18,140 genes measured in the Huang et al. study, 42% showed significant variability in expression levels due to genotype. In our study, 11% of genes are differentially expressed when comparing RAL73 to RAL 783, and 16% of genes are differentially expressed when comparing RAL306 to RAL853 (while controlling for infection status). The larger percentage found in the Huang et al. study may be attributed to their quantification of variation across 205 strains versus across two strains (per batch) in our study.

In contrast to the consistent effect of genotype, we found that the differences in transcription explained by *Wolbachia* infection were variable between the two batches. *Wolbachia* infection explained a small and non-significant proportion of variation in batch 2, but it explained approximately as much variation as genotype in batch 1. These differences in the explanatory power of infection could be due to greater transcriptional differences between samples within RAL306 and RAL853 that limit the power to detect variation due to infection status. The results could also reflect true biological differences in the degree to which *Wolbachia* affect ovarian transcription in different genotypes. A combination of these and other factors could be contributing, and future studies will be needed to disentangle the transcriptional variation due to technical versus biological differences between batches.

Though we cannot parse out the differences due to genotype versus batch across all four genotypes studied, we can compare the effect of genotype within each batch. We found a significant interaction between genotype and infection status for the samples in batch 1, suggesting the effect of *Wolbachia* on transcription depends on the genotype of the host. That a microbe’s effect depends on host genotype is increasingly documented in many systems (Ko, et al. 2009; Mateus, et al. 2019; Small, et al. 2017). The symbioses associated with *Wolbachia* vary widely by host, ranging from commensal to parasitic to mutualistic. Our finding shows that the specificity of *Wolbachia*’s relationship to its host also applies to different genotypes within the same species and that this specificity is reflected at the transcriptional level.

### Global transcription patterns reflect symbiosis between D. melanogaster and W. pipientis

We found that *Wolbachia* infection did not explain the majority of transcriptional variation among these samples. Although certain microbial infections significantly alter host transcription of many genes (Aprianto, et al. 2018; Camilios-Neto, et al. 2014; Rienksma, et al. 2019), other infections exert a more limited effect {He, 2019 #135} (Camp, et al. 2014; Rawls, et al. 2004; Small, et al. 2017). Parasitic and obligately mutualistic microbes often impose sweeping changes to host transcription. For example, a study of host tissue-specific transcriptional response to the pathogenic bacterium *Y. pseudotuberculosis* revealed substantial changes in expression. In that study1,336 genes had at least a four-fold change in expression (Nuss, et al. 2017), while our study identified only 210 and 154 genes with this extreme fold change in batch 1 and batch 2. Manipulations of obligately mutual microbes have also revealed sweeping changes to the host transcriptome (Bing, et al. 2017; Mateus, et al. 2019). While *Wolbachia* is considered an obligate mutualist in some hosts (Miller, et al. 2010; Taylor, et al. 2013) and exerts parasitic effects in other hosts (Bourtzis, et al. 1996), it does not exhibit this extreme relationship with *D. melanogaster*. The limited and variable transcriptional changes we observe is consistent with this form of symbiotic relationship.

These changes are also consistent with recent work examining the effect of *Wolbachia* infection on gene expression. In comparing whole animal transcriptomes, *Wolbachia* infection was associated with differential regulation of 285 genes that were either differentially expressed or showed significant changes in isoform use (Lindsey, et al. 2021). Similar patterns were found in ovary transcriptomic data and proteomic data{Christensen, 2016 #138;He, 2019 #135}. This is on the scale of what we observe in the current study. Genes that show altered patterns of gene expression belong to pathways associated with stress response, transcription and translation, recombination, cell cycle checkpoint (Lindsey, et al. 2021), which is similar to what we find as well.

### Genomic Location of DE genes does not support model of increased recombination via global increases in transcription

We found that genes with differential expression in *Wolbachia*-infected flies were randomly distributed across each chromosome arm. We also found that the differentially expressed genes had approximately equal levels of upregulation and downregulation. These findings are in contrast to a model in which recombination increases with *Wolbachia* infection because *Wolbachia* causes global upregulation of genes. We tested this hypothesis because across many eukaryotes, sites of recombination are correlated with sites of increased transcription (Aguilera and Gaillard 2014), and it is proposed that this correlation arises because transcription promotes recombination to resolve genomic instability (Gottipati and Helleday 2009). This correlation exists in *D. melanogaster*, as genes with ovarian transcription above 1 FPKM are concentrated in sites of known recombination (Adrian and Comeron 2012).

If the observed increase in recombination at the X-chromosome locus associated with *Wolbachia* infection (Singh 2019) was due to increased transcription in this region, then we would observe higher numbers of upregulated genes at this region. We instead observe approximately equal numbers of upregulated and downregulated genes at this region. We looked at this region in particular because we know it is associated with increased recombination under *Wolbachia* infection, but it is also possible that *Wolbachia* could increase transcription at other genomic locations we have not yet quantified. However, this trend of randomness extends to the rest of the genome: upregulated genes are distributed evenly across the genome, and we see approximately equal distribution of upregulated and downregulated genes (Figure 2). The equal distribution of differentially expressed genes across the genome and the similarity in the numbers of up- and down-regulated genes does not support the hypothesis that *Wolbachia* infection promotes increased recombination via increased transcription across many loci.

### Differentially expressed genes overlap with previous meiosis studies

We found 1,104 genes differentially regulated with *Wolbachia* infection in batch 1, and 96 genes in batch 2. We were interested in characterizing changes in gene transcription to identify candidate genes that mediate *Wolbachia*-associated plastic recombination. Previous research supports a connection between altered gene expression and altered recombination, as some recombination genes are known to affect recombination in a dosage-sensitive manner (Reynolds, et al. 2013; Ziolkowski, et al. 2017). Although genes underlying other the *Wolbachia*-induced cytoplasmic incompatibility phenotype has been identified (Beckmann, et al. 2017; LePage, et al. 2017), the genetic basis of *Wolbachia*-associated plastic recombination is unknown. Since the X-chromosome interval was previously found to exhibit increased recombination in *Wolbachia*-infected flies (Singh 2019), we compared our differentially expressed genes to a list of 111 genes significantly associated with variation in recombination at the X-chromosome locus as determined by a GWAS (Hunter, et al. 2016a). We found ten genes in common between these two studies: *Pka-C3, px, ND-23, oat, DAAM, Tgi, dpr6, spri, Msp3000* (batch 1), and *CG11200* (batch 2). These genes do not have annotated functions in meiosis, suggesting an unknown mechanism for modifying recombination rate.

In addition, we found commonalities with a list of 25 genes from a 2002 review paper of experimentally-confirmed meiosis genes (McKim, et al. 2002). Four genes reviewed in this paper were differentially expressed in our study: *c(2)m, sub, ncd* (batch 1), and *ord* (batch 2). Among the genes involved in early meiosis is *c(2)m*, which is involved in SC assembly. Since *c(2)m* suppresses crossovers (Manheim and McKim 2003), its significant downregulation in batch 1 may act to release the suppression of crossovers, ultimately resulting in more crossovers during meiosis. We also identified genes known to be involved in late meiosis, *sub* and *ncd*, which both play a role in the formation of the meiotic spindle pole. This combination of early and late meiosis genes provides an avenue of further study into the mechanisms of *Wolbachia*-induced plastic recombination.

### Lack of overlap among differentially expressed genes

Our results merit discussion particularly with regard to two previous studies looking at the effects of *Wolbachia* infection on transcription in whole flies {Lindsey, 2021 #2} and ovaries {He, 2019 #135}. Interestingly, of the genes that are identified as differentially expressed across the three studies, only two emerge as differentially expressed in all three studies: FBgn0046776 and FBgn0051619. The former is a pseudogene and the latter encodes *no long nerve cord*. This is not due to any one dataset in particular, as only nine genes overlap between the He et al. and Lindsey et al. study, thirty-five genes intersect between the current study and the whole animal Lindsey et al. study, and six genes intersect between the current study and the ovarian He et al. study.

At first blush, the lack of overlap may seem surprising. However, given the findings in all three studies that only a small fraction of the transcriptome is differentially expressed in response to infection, coupled with our observation that host genotype is the major determinant of gene expression variation, perhaps the lack of overlap is somewhat expected given that different host strains were used in the work. Moreover, other environmental factors were sure to vary as well including diet and maternal age. This finding suggests that moving forward, for the purposes of reproducibility across labs, testing for the effects of a particular treatment on gene expression should involve multiple, preferably overlapping host genotypes and highly controlled environmental factors.

### GO terms provide direction for future experiments investigating cell cycle processes

The percent of the transcriptome affected in our study (6% and 0.5%) is similar to a previous study which also did not find large-scale changes in ovarian transcription. It was recently found that 296 genes (2.2%) were differentially expressed when comparing ovaries of *D. melanogaster* infected and uninfected with *Wolbachia* (He, et al. 2019). Also consistent with this limited effect, a study of the *D. melanogaster and D. simulans* ovarian proteome found 61/549 proteins (11%) were differentially regulated in *D. melanogaster* and 49/449 (11%) were differentially regulated in *D. simulans* (Christensen, et al. 2016). Though these two studies and ours are the only ones that specifically focus on the ovarian activity of *D. melanogaster* under *Wolbachia* infection, a number of others have characterized the transcriptomic response to *Wolbachia* infection in various species and tissues. These studies confirm the effect of *Wolbachia* depends on the host, as different processes were found to be affected, including reproduction, immunity, and stress response (Brennan, et al. 2008; Chevalier, et al. 2012; Hughes, et al. 2011; Kremer, et al. 2012; Lindsey, et al. 2021; Pan, et al. 2012; Rao, et al. 2012; Xi, et al. 2008; Zheng, et al. 2011).

The differentially expressed genes from batch 1 are enriched in processes related to mitotic and meiotic cell division. Five of the top six most enriched significant GO terms are related to late cell division processes including chromosome segregation: “spindle midzone assembly”, “mitotic spindle elongation”, “kinetochore organization”, “attachment of spindle microtubules to kinetochore”, and “mitotic spindle assembly checkpoint”. Some genes in batch 2 are also associated with cell division functions, suggesting that altered cell division is consistently associated with *Wolbachia* infection in these samples. Our finding that genes associated with cell division processes are differentially regulated in *Wolbachia*-infected flies is consistent with a previous study, which found that a cell division suppressor protein 14-3-3 zeta was downregulated in *Wolbachia*-infected flies (Christensen, et al. 2016). Furthermore, the transcriptomic changes to cell division is consistent with a previously documented phenotype associated with *Wolbachia*—increased mitosis in germline stem cells associated with a four-fold increase in egg production in *D. mauritiana* (Fast, et al. 2011). Previous work on two strains used in this study, RAL306 and RAL853, also suggest that *Wolbachia* infection is associated with increased reproductive output (Singh 2019). Together, these findings merit further investigation into how *Wolbachia* influence both increased reproductive output and increased proportion of recombinant offspring.

In contrast to our study, the He et al. 2019 study did not identify any GO terms related to cell division functions. The differences in results could be related to the different genotypes used for the different studies. Although our study included multiple genotypes, it did not include the genotype studied previously (He, et al. 2019). Given that genotype has a large effect on transcription in *D. melanogaster* and that our data indicate transcriptional variation due to infection depends on host genotype, the differences between the two studies may result from differences in genotypes employed in the experiments. In addition, Wolbachia titer and phenotypes may vary as a consequence of many factors including diet, maternal age, and temperature, to name a few. Though temperature and light:dark cycle appear identical between the two studies, sufficient information is not provided to compare the effects of diet and/or maternal age. Thus, these factors may also contribute to the differences between the two studies. Future studies should address this variation by including a diversity of genetic backgrounds and environmental conditions.

### Other enriched terms are consistent with previous studies of the effect of Wolbachia infection on host transcription

We found a number of other processes enriched in *Wolbachia*-infected flies that can largely be grouped into two categories: metabolism and translation. Altered metabolism is frequently associated with host-microbe interactions (Maslowski 2019), and our findings fit this pattern. *Wolbachia* infection is associated with depletion of certain classes of lipids in the mosquito *Aedes albopictus* (Molloy, et al. 2016), increased host resistance to iron depletion (Brownlie, et al. 2009), and altered host production of dopamine-dependent arylalkylamine N-acetyltransferase (Gruntenko, et al. 2017). We also found altered N-acetylneuraminate catabolism associated with *Wolbachia* infection in our study, along with other metabolic processes such as “negative regulation of insulin secretion,”, “glucosamine catabolic process,” “glutathione metabolic process,” and “chitin metabolic process.” Altered metabolism was found in the two other studies of *Wolbachia*-infected ovaries of *D. melanogaster*: Christenson et al. (2016) found differential levels of proteins related to carbohydrate transfer and metabolism, and He et al. (2019) find differential regulation of starch and sucrose metabolism genes.

Apart from altered cell division, we found the processes most affected by *Wolbachia* infection are those related to translation, protein synthesis, and ribosome synthesis. Furthermore, most of these differentially expressed genes are downregulated. These results are consistent with previous work (Christensen, et al. 2016), who also found downregulation of protein synthesis proteins in both *Wolbachia*-infected *D. melanogaster* and *D. simulans*. Interestingly, a recent study found that perturbing assembly of translation components through RNAi results in increased *Wolbachia* titer, further supporting a link between decreased translation and *Wolbachia* infection (Grobler, et al. 2018).

## Conclusions

We investigated how ovarian transcription is altered with *Wolbachia* infection in this interesting host-microbe system, particularly with respect to increased recombination in the host. We found that *Wolbachia* infection contributes in a limited and variable way to host ovarian transcription, and demonstrated for the first time that *Wolbachia*’s effect on ovarian transcription is mediated by the host’s genotype. Altered ovarian cell division processes are associated with *Wolbachia* infection and identified a set of candidate genes that will allow further exploration of the link between *Wolbachia* infection and increased recombination. The downregulation of *c(2)m* in *Wolbachia*-infected flies, along with its known role as a suppressor of crossovers (71), make this gene a promising candidate for future studies. Other genes of interest include those involved in spindle and kinetochore processes (e.g. *sub, polo, incenp*), which are highly enriched terms in *Wolbachia*-infected flies. These identified transcriptomic changes will guide future studies of *Wolbachia*’s symbiosis with its *D. melanogaster* host and its relation to plastic recombination.

## Methods

### Drosophila stocks

We used four strains from the Drosophila Genome Reference Panel (DRGP) (Mackay, et al. 2012) for this study: RAL73, RAL783, RAL306, and RAL853. These lines have standard chromosome arrangements and are naturally infected with *Wolbachia pipientis* (Huang, et al. 2015). Genomic analysis confirmed that these flies are infected with *w*Mel haplotype 1 (Richardson, et al. 2012). To generate *Wolbachia*-uninfected flies, we raised these strains on food containing tetracycline at a concentration of 0.25 mg/mL media for two generations. To reintroduce the natural microbiome without *Wolbachia, Wolbachia*-uninfected flies were subsequently raised on standard food for at least ten additional generations before transcriptome data were obtained (Singh 2019). *Wolbachia* infection status was confirmed prior to experimentation using a PCR-based assay with *Wolbachia*-specific primers targeted to *Wolbachia*-specific protein (Jeyaprakash and Hoy 2000). *Wolbachia* infection status was also confirmed for each sample after RNA-sequencing by testing for reads that aligned successfully to the *Wolbachia* reference genome. Throughout their rearing, flies were kept at 25 degrees Celsius on a 12:12 hour light:dark cycle.

### Ovary dissections and sequencing

We placed carbon dioxide-anesthetized three-day-old virgin females in phosphate-buffer saline, dissected their ovaries, and froze the ovaries on dry ice. We prepared 4 replicates of each of the 4 genotypes for each infection status, with 10 pairs of pooled ovaries comprising each replicate. Dissections, library preparations, and sequencing were performed in two batches two months apart: all dissections on the RAL783 and RAL73 samples were performed on one day, and all dissections of RAL306 and RAL853 were performed in one day two months later. Each replicate of 10 pairs of ovaries remained frozen until total RNA was isolated from them using the Zymo Direct-zol kit. Integrity of the isolated RNA was evaluated using an Agilent Fragment Analyzer, and RQN values ranged from 8.9-10. Poly-A-selected mRNA-seq libraries were prepared using the NuGen (Tecan) Universal Plus kit, which introduces chemical fragmentation of the mRNA prior to first strand synthesis. RNA isolations and RNA-seq libraries were prepared and sequenced at the University of Oregon Genomics and Cell Characterization Core Facility. Libraries were sequenced on a HiSeq 4000 using single end, 150-bp reads.

### Sequence processing and alignment

Raw reads were aligned to the *Drosophila melanogaster* reference genome from the Berkeley Drosophila Genome Project (BDGP) assembly release 6 with STAR aligner v. 2.5.3a (Dobin, et al. 2013) with default parameters. Batch 1 (comprised of strains RAL73 and RAL783) generated an average of 22,443,144 reads with 98% aligning uniquely to the reference genome, and batch 2 (comprised of RAL306 and RAL853) generated on average 21,936,370 reads with 89% aligning uniquely to the reference genome (Supplementary Table 1).

### Differential gene expression analysis

All analyses were conducted using R (v. 3.4.3) packages. For each sample we counted the number of uniquely mapped reads using GenomicAlignments (v. 3.6-intel-2017b) (Lawrence, et al. 2013) with default parameters. We then used DESeq2 (Love, et al. 2014) to perform normalization and differential expression analysis on gene count data. Specifically, we fit a model of expression level for each gene as determined by the effect of infection status, while controlling for genotype and the interaction of infection status and genotype. DESeq2 uses a model particularly suited for the overdispersion of RNA-seq data based on the negative binomial distribution. We set the uninfected samples as the base level, so “upregulation” refers to higher normalized read counts in the infected flies. DESeq2 uses the Benjamini and Hochberg (Hochberg and Benjamini 1990) method by default to control the false discovery rate (FDR). We define differentially expressed genes after controlling the FDR at 0.05. We did not enforce a fold-change threshold in addition to statistical significance based on the premise that genes in the recombination pathway are precisely regulated so that even a small change in expression levels could reflect a functional change in meiosis (Reynolds, et al. 2013; Ziolkowski, et al. 2017). Using the model above, we conducted gene-wise differential expression tests separately for libraries from batch 1 and batch 2, because genotypes were not replicated across batches. Given this design property, and the clearly strong transcriptome-wide batch effect (Fig 1), performing the analyses separately for each batch allowed us to properly evaluate effects of genotype (and genotype-by-*Wolbachia* interaction) on gene expression independently of batch.

### Multivariate Analysis with nMDS and perMANOVA

To visualize differences in expression among 17,377 transcripts across all samples, we used Bray-Curtis dissimilarity to generate overall and batch-specific nMDS ordination plots. We quantified the percentage of dissimilarity explained by infection status, genotype, and the interaction of infection and genotype with a permutational multivariate analysis of variance (perMANOVA) test. Similar to the gene-wise differential expression analyses above, we accounted for batch by performing the perMANOVA test separately on each of the two batches and accounted for genotype within each test by limiting permutations to samples within genotypes. We used the R package vegan v. 2.5-5, (Oksanen, et al. 2007) to calculate distances, plot nMDS, and perform the perMANOVA test.

### Outlier Ordination

Initial visualization with an nMDS plot revealed that sample RAL73 *Wolbachia*-infected A (RAL73 w+A) differed from the other three replicates of RAL73 infected samples, as well as all other samples in this study. To determine whether this sample represented an outlier with significant influence on the results of the differential expression analysis, we examined Cook’s distance on the normalized count data, as calculated by DESeq2. A higher Cook’s distance for a sample indicates more leverage in determining a given transcript’s fold change. We compared the average Cook’s distance for each sample across all transcripts and the number of times each sample contained the highest Cook’s distance for each transcript (Supplementary Figure 1). Replicate samples should not differ much in their distribution of Cook’s distances, but we found that the average Cook’s distance for sample RAL73w+ replicate A was much higher than all other samples and most often contains the maximum Cook’s distance across all transcripts. These results suggest sample RAL73w+A has aberrant expression of many genes. Although DESeq2 can manage outliers for transcripts of genes if there are replicates of each factor, we decided that a sample which consistently contains an outlier across many genes likely represents not true biological signal but rather an error of preparation. This distribution of Cook’s distances led us to exclude RAL73w+A from all subsequent analyses.

### Gene Ontology Analysis

We identified Gene Ontology (GO) descriptions for the differentially expressed genes using the Panther biological process classification system, as implemented in Panther’s data mapping online tool (Thomas, et al. 2003). Panther classifies each GO term into nested levels, and we report the level of nesting for GO terms in order to provide more specific information. We then tested whether any GO terms were overrepresented in each group of differentially expressed genes using Panther’s statistical overrepresentation test, which uses a Fisher’s exact test with False Discovery Rate (FDR) controlled at 0.1 and all annotated genes in the *Drosophila melanogaster* genome as reference.

### Distribution of DE Genes Across Genome

To test whether differentially expressed genes are randomly distributed across the genome, we performed a non-parametric run’s test for randomness (Bradley 1968). We categorized each gene as differentially expressed or not, based on the same criteria of FDR controlled at 0.05. We then placed each gene’s differential expression information in order of placement across the genome, using the gene start site as the location. A switch from differentially expressed to not, or vice-versa, was considered a complete run. We visualized these patterns of gene expression across the genome with the R package karyoploteR (Gel and Serra 2017) version 1.8.8. For each chromosome, we depict the differentially expressed genes at their location in the BDGP6 *D. melanogaster* reference genome.

## Supporting information

AdditionalFile1

AdditionalFile2

## Declarations

### Ethics approval and consent to participate

Not applicable.

### Consent for publication

Not applicable.

### Availability of data and materials

The data supporting the results of this article will be available at the NCBI SRA repository upon publication.

### Authors’ contributions

SF performed all analyses and wrote the manuscript. NDS designed the experiment and advised with analyses and manuscript preparation. WAC and CMS advised with analyses and manuscript preparation. Any correspondence and request for materials should be addressed to NDS. All authors read and approved the final manuscript.

## Acknowledgements

The authors would like to thank the members of the Singh and Cresko labs for their valuable feedback on this manuscript.

## Literature Cited

Adrian A, Comeron JM. 2012. The Drosophila early ovarian transcriptome provides insight into the molecular causes of recombination rate variation. Poster presentation at the Annual Drosophila Meeting; Washington. DC.

Aggarwal DD, et al. 2019. Desiccation-induced changes in recombination rate and crossover interference in Drosophila melanogaster: evidence for fitness-dependent plasticity. Genetica 147: 291–302. doi: 10.1007/s10709-019-00070-6

Agrawal AF, Hadany L, Otto SP 2005. The evolution of plastic recombination. Genetics 171: 803–812. doi: Doi 10.1534/Genetics.105.041301

Aguilera A, Gaillard H 2014. Transcription and Recombination: When RNA Meets DNA. Cold Spring Harbor Perspectives in Biology 6. doi: ARTN a01654310.1101/cshperspect.a016543

Andronic L 2012. Viruses as triggers of DNA rearrangements in host plants. Canadian Journal of Plant Science 92: 1083–1091. doi: 10.4141/cjps2011-197

Aprianto R, Slager J, Holsappel S, Veening JW 2018. High-resolution analysis of the pneumococcal transcriptome under a wide range of infection-relevant conditions. Nucleic Acids Research 46: 9990–10006. doi: 10.1093/nar/gky750

Ballard JWO 2004. Sequential evolution of a symbiont inferred from the host: Wolbachia and Drosophila simulans. Molecular Biology and Evolution 21: 428–442.

Beckmann JF, Ronau JA, Hochstrasser M 2017. A Wolbachia deubiquitylating enzyme induces cytoplasmic incompatibility. Nature Microbiology 2. doi: ARTN 1700710.1038/nmicrobiol.2017.7

Belyaev DK, Borodin PM 1982. The Influence of Stress on Variation and Its Role in Evolution. Biologisches Zentralblatt 101: 705–714.

Bing XL, et al. 2017. Unravelling the relationship between the tsetse fly and its obligate symbiont Wigglesworthia: transcriptomic and metabolomic landscapes reveal highly integrated physiological networks. Proceedings of the Royal Society B-Biological Sciences 284. doi: ARTN 2017036010.1098/rspb.2017.0360

Bourtzis K, Nirgianaki A, Markakis G, Savakis C 1996. Wolbachia infection and cytoplasmic incompatibility in Drosophila species. Genetics 144: 1063–1073.

Bradley JV. 1968. Distribution-free statistical tests. Englewood Cliffs, N.J.,: Prentice-Hall.

Bradshaw AD 1965. Evolutionary Significance of Phenotypic Plasticity in Plants. Advances in Genetics 13: 115–155.

Brakefield PM 1987. Tropical Dry and Wet Season Polyphenism in the Butterfly Melanitis-Leda (Satyrinae) - Phenotypic Plasticity and Climatic Correlates. Biological Journal of the Linnean Society 31: 175–191. doi: DOI 10.1111/j.1095-8312.1987.tb01988.x

Brem RB, Yvert G, Clinton R, Kruglyak L 2002. Genetic dissection of transcriptional regulation in budding yeast. Science 296: 752–755. doi: 10.1126/science.1069516

Brennan LJ, Keddie BA, Braig HR, Harris HL 2008. The Endosymbiont Wolbachia pipientis Induces the Expression of Host Antioxidant Proteins in an Aedes albopictus Cell Line. Plos One 3. doi: ARTN e208310.1371/journal.pone.0002083

Brownlie JC, et al. 2009. Evidence for Metabolic Provisioning by a Common Invertebrate Endosymbiont, Wolbachia pipientis, during Periods of Nutritional Stress. Plos Pathogens 5. doi: ARTN e100036810.1371/journal.ppat.1000368

Bryant KN, Newton ILG 2020. The Intracellular Symbiont Wolbachia pipientis Enhances Recombination in a Dose-Dependent Manner. Insects 11. doi: 10.3390/insects11050284

Camilios-Neto D, et al. 2014. Dual RNA-seq transcriptional analysis of wheat roots colonized by Azospirillum brasilense reveals up-regulation of nutrient acquisition and cell cycle genes. Bmc Genomics 15. doi: ARTN 37810.1186/1471-2164-15-378

Camp JG, et al. 2014. Microbiota modulate transcription in the intestinal epithelium without remodeling the accessible chromatin landscape. Genome Research 24: 1504–1516. doi: 10.1101/gr.165845.113

Chevalier F, et al. 2012. Feminizing Wolbachia: a transcriptomics approach with insights on the immune response genes in Armadillidium vulgare. Bmc Microbiology 12. doi: ARTN S110.1186/1471-2180-12-S1-S1

Christensen S, et al. 2016. Wolbachia Endosymbionts Modify Drosophila Ovary Protein Levels in a Context-Dependent Manner. Applied and Environmental Microbiology 82: 5354–5363. doi: 10.1128/Aem.01255-16

Dixon AL, et al. 2007. A genome-wide association study of global gene expression. Nature Genetics 39: 1202–1207. doi: 10.1038/ng2109

Dobin A, et al. 2013. STAR: ultrafast universal RNA-seq aligner. Bioinformatics 29: 15–21. doi: 10.1093/bioinformatics/bts635

Fast EM, et al. 2011. Wolbachia Enhance Drosophila Stem Cell Proliferation and Target the Germline Stem Cell Niche. Science 334: 990–992. doi: 10.1126/science.1209609

Fry AJ, Palmer MR, Rand DM 2004. Variable fitness effects of Wolbachia infection in Drosophila melanogaster. Heredity 93: 379–389. doi: 10.1038/sj.hdy.6800514

Fry AJ, Rand DM 2002. Wolbachia interactions that determine Drosophila melanogaster survival. Evolution 56: 1976–1981.

Gao J, Davidson MK, Wahls WP 2008. Distinct regions of ATF/CREB proteins Atf1 and Pcr1 control recombination hotspot ade6-M26 and the osmotic stress response. Nucleic Acids Research 36: 2838–2851. doi: 10.1093/nar/gkn037

Gel B, Serra E 2017. karyoploteR: an R/Bioconductor package to plot customizable genomes displaying arbitrary data. Bioinformatics 33: 3088–3090. doi: 10.1093/bioinformatics/btx346

Gottipati P, Helleday T 2009. Transcription-associated recombination in eukaryotes: link between transcription, replication and recombination. Mutagenesis 24: 203–210. doi: 10.1093/mutage/gen072

Grell RF 1978. Comparison of Heat and Inter-Chromosomal Effects on Recombination and Interference in Drosophila-Melanogaster. Genetics 89: 65–77.

Grobler Y, et al. 2018. Whole genome screen reveals a novel relationship between Wolbachia levels and Drosophila host translation. Plos Pathogens 14. doi: ARTN e100744510.1371/journal.ppat.1007445

Gruntenko NE, et al. 2017. Various Wolbachia genotypes differently influence host Drosophila dopamine metabolism and survival under heat stress conditions. BMC Evolutionary Biology 17. doi: ARTN 25210.1186/s12862-017-1104-y

Hayman DL, Parsons PA 1961. The effect of temperature, age and an inversion on recombination values and interference in the X-chromosome of Drosophila melanogaster. Genetica 32: 74–88. doi: 10.1007/BF01816087

He Z, et al. 2019. How do Wolbachia modify the Drosophila ovary? New evidences support the “titration-restitution” model for the mechanisms of Wolbachia-induced CI. Bmc Genomics 20. doi: ARTN 60810.1186/s12864-019-5977-6

Hilgenboecker K, Hammerstein P, Schlattmann P, Telschow A, Werren JH 2008. How many species are infected with Wolbachia? -a statistical analysis of current data. FEMS Microbiology Letters 281: 215–220. doi: Doi 10.1111/J.1574-6968.2008.01110.X

Hochberg Y, Benjamini Y 1990. More powerful procedures for multiple significance testing. Stat Med 9: 811–818. doi: 10.1002/sim.4780090710

Hoffmann AA, Hercus M, Dagher H 1998. Population dynamics of the Wolbachia infection causing cytoplasmic incompatibility in Drosophila melanogaster. Genetics 148: 221–231.

Hosokawa T, Koga R, Kikuchi Y, Meng XY, Fukatsu T 2010. Wolbachia as a bacteriocyte-associated nutritional mutualist. Proceedings of the National Academy of Sciences of the United States of America 107: 769–774. doi: 10.1073/pnas.0911476107

Huang W, et al. 2015. Genetic basis of transcriptome diversity in Drosophila melanogaster. Proceedings of the National Academy of Sciences of the United States of America 112: E6010–E6019. doi: 10.1073/pnas.1519159112

Hughes GL, et al. 2011. Wolbachia Infections in Anopheles gambiae Cells: Transcriptomic Characterization of a Novel Host-Symbiont Interaction. Plos Pathogens 7. doi: ARTN e100129610.1371/journal.ppat.1001296

Hunter CM, Huang W, Mackay TFC, Singh ND 2016a. The Genetic Architecture of Natural Variation in Recombination Rate in Drosophila melanogaster. Plos Genetics 12. doi: ARTN e100595110.1371/journal.pgen.1005951

Hunter CM, Robinson MC, Aylor DL, Singh ND 2016b. Genetic Background, Maternal Age, and Interaction Effects Mediate Rates of Crossing Over in Drosophila melanogaster Females. G3-Genes Genomes Genetics 6: 1409–1416. doi: 10.1534/g3.116.027631

Hussin J, Roy-Gagnon M-Hln, Gendron R, Andelfinger G, Awadalla P 2011. Age-Dependent Recombination Rates in Human Pedigrees. PLoS Genet 7: e1002251.

Hutter S, Saminadin-Peter SS, Stephan W, Parsch J 2008. Gene expression variation in African and European populations of Drosophila melanogaster. Genome Biology 9. doi: ARTN R1210.1186/gb-2008-9-1-r12

Jackson S, Nielsen DM, Singh ND 2015. Increased exposure to acute thermal stress is associated with a non-linear increase in recombination frequency and an independent linear decrease in fitness in Drosophila. BMC Evolutionary Biology 15. doi: ARTN 17510.1186/S12862-015-0452-8

Jeyaprakash A, Hoy MA 2000. Long PCR improves Wolbachia DNA amplification: wsp sequences found in 76% of sixty-three arthropod species. Insect Molecular Biology 9: 393–405. doi: Doi 10.1046/J.1365-2583.2000.00203.X

Ko DC, et al. 2009. A Genome-wide In Vitro Bacterial-Infection Screen Reveals Human Variation in the Host Response Associated with Inflammatory Disease. American Journal of Human Genetics 85: 214–227. doi: 10.1016/j.ajhg.2009.07.012

Kohl KP, Singh ND 2018. Experimental Evolution Across Different Thermal Regimes Yields Genetic Divergence in Recombination Fraction But No Divergence in Temperature-Associated Plastic Recombination. Evolution 72: 989–999.

Kon N, Schroeder SC, Krawchuk MD, Wahls WP 1998. Regulation of the Mts1-Mts2-dependent ade6-M26 meiotic recombination hot spot and developmental decisions by the Spc1 mitogen-activated protein kinase of fission yeast. Molecular and Cellular Biology 18: 7575–7583. doi: Doi 10.1128/Mcb.18.12.7575

Kremer N, et al. 2012. Influence of Wolbachia on host gene expression in an obligatory symbiosis. Bmc Microbiology 12. doi: ARTN S710.1186/1471-2180-12-S1-S7

Kriesner P, Hoffmann AA, Lee SF, Turelli M, Weeks AR 2013. Rapid Sequential Spread of Two Wolbachia Variants in Drosophila simulans. Plos Pathogens 9. doi: ARTN e100360710.1371/journal.ppat.1003607

Lawrence M, et al. 2013. Software for computing and annotating genomic ranges. PLoS Comput Biol 9: e1003118. doi: 10.1371/journal.pcbi.1003118

LePage DP, et al. 2017. Prophage WO genes recapitulate and enhance Wolbachia-induced cytoplasmic incompatibility. Nature 543: 243-+. doi: 10.1038/nature21391

Lindsey ARI, et al. 2021. Wolbachia and Virus Alter the Host Transcriptome at the Interface of Nucleotide Metabolism Pathways. mBio 12: e03472–03420.

Love MI, Huber W, Anders S 2014. Moderated estimation of fold change and dispersion for RNA-seq data with DESeq2. Genome Biology 15: 550. doi: 10.1186/s13059-014-0550-8

Mackay TFC, et al. 2012. The Drosophila melanogaster Genetic Reference Panel. Nature 482: 173–178. doi: http://www.nature.com/nature/journal/v482/n7384/abs/nature10811.html#supplementary-information

Manheim EA, McKim KS 2003. The synaptonemal complex component C(2)M regulates meiotic crossing over in Drosophila. Current Biology 13: 276–285. doi: Pii S0960-9822(03)00050-2Doi 10.1016/S0960-9822(03)00050-2

Maslowski KM 2019. Metabolism at the centre of the host-microbe relationship. Clinical and Experimental Immunology 197: 193–204. doi: 10.1111/cei.13329

Mateus ID, et al. 2019. Dual RNA-seq reveals large-scale non-conserved genotype x genotype-specific genetic reprograming and molecular crosstalk in the mycorrhizal symbiosis. Isme Journal 13: 1226–1238. doi: 10.1038/s41396-018-0342-3

McKim KS, Jang JK, Manheim EA 2002. Meiotic recombination and chromosome segregation in Drosophila females. Annual Review of Genetics 36: 205–232. doi: 10.1146/annurev.genet.36.041102.113929

Mi HY, et al. 2010. PANTHER version 7: improved phylogenetic trees, orthologs and collaboration with the Gene Ontology Consortium. Nucleic Acids Research 38: D204–D210. doi: 10.1093/nar/gkp1019

Miller WJ, Ehrman L, Schneider D 2010. Infectious Speciation Revisited: Impact of Symbiont-Depletion on Female Fitness and Mating Behavior of Drosophila paulistorum. Plos Pathogens 6. doi: ARTN e100121410.1371/journal.ppat.1001214

Modliszewski JL, Copenhaver GP 2017. Meiotic recombination gets stressed out: CO frequency is plastic under pressure. Current Opinion in Plant Biology 36: 95–102. doi: 10.1016/j.pbi.2016.11.019

Molloy JC, Sommer U, Viant MR, Sinkins SP 2016. Wolbachia Modulates Lipid Metabolism in Aedes albopictus Mosquito Cells. Applied and Environmental Microbiology 82: 3109–3120. doi: 10.1128/Aem.00275-16

Neel JV 1941. A relation between larval nutrition and the frequency of crossing over in the third chromosome of Drosophila melanogaster. Genetics 26: 506–516.

Nuss AM, et al. 2017. Tissue dual RNA-seq allows fast discovery of infection-specific functions and riboregulators shaping host-pathogen transcriptomes. Proceedings of the National Academy of Sciences of the United States of America 114: E791–E800. doi: 10.1073/pnas.1613405114

Oksanen J, et al. 2007. The vegan package. Community Ecology Package 10: 719.

Pan XL, et al. 2012. Wolbachia induces reactive oxygen species (ROS)-dependent activation of the Toll pathway to control dengue virus in the mosquito Aedes aegypti. Proceedings of the National Academy of Sciences of the United States of America 109: E23–E31. doi: 10.1073/pnas.1116932108

Plough HH 1917. The effect of temperature on crossing over in Drosophila. Journal of Experimental Zoology 24: 147–209.

Plough HH 1921. Further studies on the effect of temperature on crossing over. Journal of Experimental Zoology 32: 187–202.

Poelstra JW, et al. 2014. The genomic landscape underlying phenotypic integrity in the face of gene flow in crows. Science 344: 1410–1414. doi: 10.1126/science.1253226

Rao RU, et al. 2012. Effects of doxycycline on gene expression in Wolbachia and Brugia malayi adult female worms in vivo. J Biomed Sci 19: 21. doi: 10.1186/1423-0127-19-21

Rawls JF, Samuel BS, Gordon JI 2004. Gnotobiotic zebrafish reveal evolutionarily conserved responses to the gut microbiota. Proceedings of the National Academy of Sciences of the United States of America 101: 4596–4601. doi: 10.1073/pnas.0400706101

Reynolds A, et al. 2013. RNF212 is a dosage-sensitive regulator of crossing-over during mammalian meiosis. Nature Genetics 45: 269–278. doi: 10.1038/ng.2541

Richardson MF, et al. 2012. Population Genomics of the Wolbachia Endosymbiont in Drosophila melanogaster. PLoS Genetics 8. doi: ARTN E1003129 Doi 10.1371/Journal.Pgen.1003129

Rienksma RA, Schaap PJ, dos Santos VAPM, Suarez-Diez M 2019. Modeling Host-Pathogen Interaction to Elucidate the Metabolic Drug Response of Intracellular Mycobacterium tuberculosis. Frontiers in Cellular and Infection Microbiology 9. doi: ARTN 144 10.3389/fcimb.2019.00144

Scheepens JF, Deng Y, Bossdorf O 2018. Phenotypic plasticity in response to temperature fluctuations is genetically variable, and relates to climatic variability of origin, in Arabidopsis thaliana. Aob Plants 10. doi: ARTN ply043 10.1093/aobpla/ply043

Serga S, Maistrenko O, Rozhok A, Kozeretska I 2014. Fecundity as one of possible factors contributing to the dominance of the wMel genotype of Wolbachia in natural populations of Drosophila melanogaster. Febs Journal 281: 741–741.

Singh ND 2019. Wolbachia Infection Associated with Increased Recombination in Drosophila. G3-Genes Genomes Genetics 9: 229–237. doi: 10.1534/g3.118.200827

Singh ND, et al. 2015. Fruit flies diversify their offspring in response to parasite infection. Science 349: 747–750. doi: 10.1126/science.aab1768

Small CM, Milligan-Myhre K, Bassham S, Guillemin K, Cresko WA 2017. Host Genotype and Microbiota Contribute Asymmetrically to Transcriptional Variation in the Threespine Stickleback Gut. Genome Biology and Evolution 9: 504–520. doi: 10.1093/gbe/evx014

Solignac M, Vautrin D, Rousset F 1994. Widespread Occurrence of the Proteobacteria Wolbachia and Partial Cytoplasmic Incompatibility in Drosophila-Melanogaster. Comptes Rendus De L Academie Des Sciences Serie Iii-Sciences De La Vie-Life Sciences 317: 461–470.

Stapley J, Feulner PGD, Johnston SE, Santure AW, Smadja CM 2017. Variation in recombination frequency and distribution across eukaryotes: patterns and processes. Philosophical Transactions of the Royal Society B-Biological Sciences 372. doi: ARTN 20160455 10.1098/rstb.2016.0455

Stern C 1926. An effect of temperature and age on crossing over in the first chromosome of Drosophila melanogaster. Proceedings of the National Academy of Sciences of the United States of America 12: 530–532.

Taylor MJ, Voronin D, Johnston KL, Ford L 2013. Wolbachia filarial interactions. Cellular Microbiology 15: 520–526. doi: 10.1111/cmi.12084

Thomas PD, et al. 2003. PANTHER: A library of protein families and subfamilies indexed by function. Genome Research 13: 2129–2141. doi: 10.1101/gr.772403

Tollrian R 1995. Predator-Induced Morphological Defenses - Costs, Life-History Shifts, and Maternal Effects in Daphnia-Pulex. Ecology 76: 1691–1705. doi: Doi 10.2307/1940703

Vavre F, Girin C, Bouletreau M 1999. Phylogenetic status of a fecundity-enhancing Wolbachia that does not induce thelytoky in Trichogramma. Insect Molecular Biology 8: 67–72. doi: DOI 10.1046/j.1365-2583.1999.810067.x

Verbon EH, Liberman LM 2016. Beneficial Microbes Affect Endogenous Mechanisms Controlling Root Development. Trends in Plant Science 21: 218–229. doi: 10.1016/j.tplants.2016.01.013

Weeks AR, Turelli M, Harcombe WR, Reynolds KT, Hoffmann AA 2007. From parasite to mutualist: Rapid evolution of Wolbachia in natural populations of Drosophila. Plos Biology 5: 997–1005. doi: ARTN e114 10.1371/journal.pbio.0050114

Werren JH, Baldo L, Clark ME 2008. Wolbachia: master manipulators of invertebrate biology. Nature Reviews Microbiology 6: 741–751. doi: 10.1038/nrmicro1969

Xi ZY, Gavotte L, Xie Y, Dobson SL 2008. Genome-wide analysis of the interaction between the endosymbiotic bacterium Wolbachia and its Drosophila host. Bmc Genomics 9. doi: ARTN 1 10.1186/1471-2164-9-1

Zheng Y, et al. 2011. Differentially expressed profiles in the larval testes of Wolbachia infected and uninfected Drosophila. Bmc Genomics 12. doi: ARTN 595 10.1186/1471-2164-12-595

Zilio G, Moesch L, Bovet N, Sarr A, Koella JC 2018. The effect of parasite infection on the recombination rate of the mosquito Aedes aegypti. Plos One 13. doi: ARTN e0203481 10.1371/journal.pone.0203481

Ziolkowski PA, et al. 2017. Natural variation and dosage of the HEI10 meiotic E3 ligase control Arabidopsis crossover recombination. Genes & Development 31: 306–317. doi: 10.1101/gad.295501.116

Zug R, Hammerstein P 2012. Still a Host of Hosts for Wolbachia: Analysis of Recent Data Suggests That 40% of Terrestrial Arthropod Species Are Infected. Plos One 7. doi: ARTN e38544 DOI 10.1371/journal.pone.0038544

