## AdditionalFile1 for "Candidate Genes Underlying *Wolbachia*-Associated Plastic Recombination Revealed by Ovarian Transcriptomics of *D. melanogaster*"

**<u>Electronic Supplementary Material</u>**

**Additional file 1:** Supplementary Figure 1: Outlier investigation through metrics of Cook's distance. Supplementary Table 1: Sequencing Statistics. (.pdf)

**Additional file 2:** Supplementary Table 2: Differentially expressed genes identified in batch 1. Supplementary Table 3: Differentially expressed genes identified in batch 2. Supplementary Table 4: Differentially expressed genes common to both batch 1 and batch 2. Supplementary Table 5. Overrepresented GO terms associated with differentially expressed genes from batch 1. Supplementary Table 6. Overrepresented GO terms associated with differentially expressed genes from batch 2. (.xlsx)

Additional File 1

**a)**

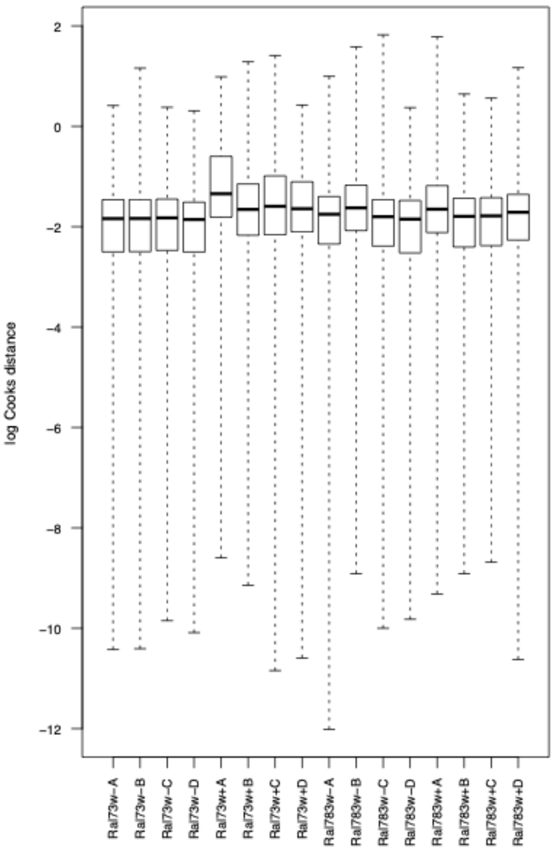


**b)**

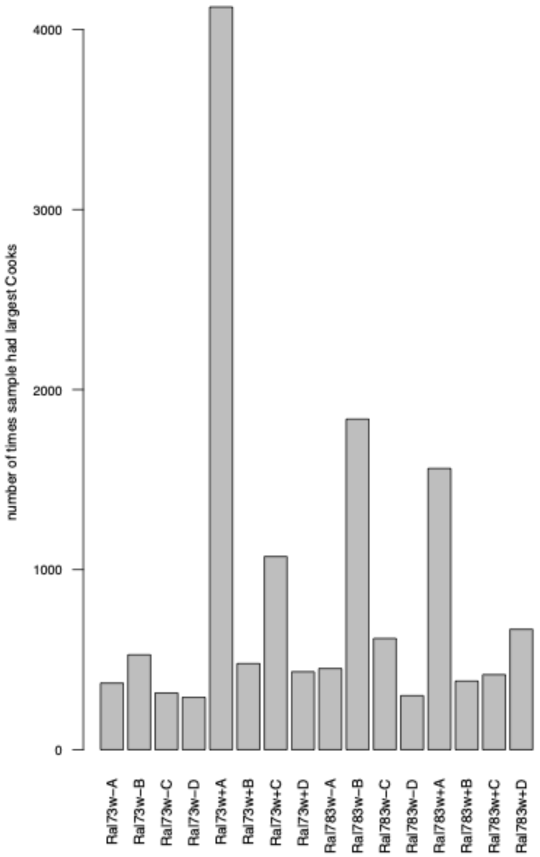

**Supplementary Figure 1:** Outlier investigation through metrics of Cook's distance. **a)** For every transcript with count data, Cook's distance was measured for each sample. The distribution of log Cook's distances for each sample is shown. RAL73w+ has higher average Cooks distance than all other samples. **b)** The number of times each sample had the highest Cook's distance for a given transcript is plotted. RAL73w+A most often has the highest Cook's distance.

| Batch 1 | | | Batch 2 | | |
|---|---|---|---|---|---|
| sample ID | # reads mapped | %reads mapped | sample ID | # reads mapped | %reads mapped |
| RAL73w - A | 25,580,060 | 98.21 | RAL306w - A | 30,181,092 | 89.9 |
| RAL73w - B | 23,340,083 | 97.67 | RAL306w - B | 21,532,464 | 88.6 |
| RAL73w - C | 23,830,990 | 98.1 | RAL306w - C | 29,151,147 | 89.87 |
| RAL73w - D | 23,886,917 | 98.18 | RAL306w - D | 24,077,440 | 86.97 |
| RAL73w + A | 23,461,064 | 98.14 | RAL306w + A | 25,333,773 | 90.52 |
| RAL73w + B | 22,213,114 | 98.15 | RAL306w + B | 23,015,730 | 90.09 |
| RAL73w + C | 22,414,516 | 98.25 | RAL306w + C | 21,057,850 | 90.57 |
| RAL73w + D | 20,165,444 | 98.2 | RAL306w + D | 19,633,517 | 90.35 |
| RAL783w - A | 24,508,882 | 98.29 | RAL853w - A | 20,800,548 | 90.93 |
| RAL783w - B | 21,975,849 | 97.75 | RAL853w - B | 16,410,190 | 90.76 |
| RAL783w - C | 20,906,567 | 98.09 | RAL853w - C | 22,790,031 | 87.11 |
| RAL783w - D | 22,989,934 | 98.2 | RAL853w - D | 18,623,251 | 89.76 |
| RAL783w + A | 20,287,678 | 98.22 | RAL853w + A | 23,365,471 | 90.6 |
| RAL783w + B | 21,266,231 | 98.04 | RAL853w + B | 15,723,826 | 89.15 |
| RAL783w + C | 21,347,772 | 98.01 | RAL853w + C | 22,380,358 | 88.8 |
| RAL783w + D | 20,915,208 | 97.94 | RAL853w + D | 16,905,243 | 90.06 |
| | mean 22,443,144 | mean 98.09 | | mean 21,936,370 | mean 89.63 |

**Supplementary Table 1:** Sequencing Statistics. Shown are the sample ID for batch 1 and batch 2, the total number of reads that mapped uniquely to the genome for each sample, and the percent of total reads that mapped uniquely to the genome. The mean number and percentage across samples in each batch are also shown.
